# Alzheimer cells on their way to derailment show selective changes in protein quality control network

**DOI:** 10.1101/2020.05.17.099465

**Authors:** Margreet B. Koopman, Stefan G.D Rüdiger

## Abstract

Alzheimer’s Disease is driven by protein aggregation and is characterised by accumulation of Tau protein into neurofibrillary tangles. In healthy neurons the cellular protein quality control is successfully in charge of protein folding, which raises the question to which extent this control is disturbed in disease. Here we describe that brain cells in Alzheimer’s Disease show very specific derailment of the protein quality control network. We performed a meta-analysis on the Alzheimer’s Disease Proteasome database, which provides a quantitative assessment of disease-related proteome changes in six brain regions in comparison with age-matched controls. We noted that levels of all paralogues of the conserved Hsp90 chaperone family are reduced, while most other chaperones – or their regulatory co-chaperones – do not change in disease. The notable exception is a select group consisting of the stress inducible HSP70, its nucleotide exchange factor BAG3 – which links the Hsp70 system to autophagy – and neuronal small heat shock proteins, which are upregulated in disease. They are all members of a cascade controlled in the stress response, channelling proteins towards a pathway of chaperone assisted selective autophagy. Together, our analysis reveals that in an Alzheimer’s brain, with exception of Hsp90, the players of the protein quality control are still present in full strength, even in brain regions most severely affected in disease. The specific upregulation of small heat shock proteins and HSP70:BAG3, ubiquitous in all brain areas analysed, may represent a last, unsuccessful attempt to advert neuronal cell death.

## INTRODUCTION

Neurodegenerative diseases are a group of diseases characterized by progressive neuronal degeneration, of which Alzheimer’s Disease (AD) is the most prominent one. Symptoms of AD include severe memory loss and cognitive decline and are often accompanied by changes in personality (Bature et al., 2017). On molecular level, AD shows the accumulation of two distinct proteins; extracellular plaques of amyloid-ß and intracellular formation of neurofibrillary tangles of Tau; a microtubule associated protein assisting in microtubule stability and regulating axonal transport (Drubin and Kirschner, 1986; Goedert et al., 2017). Under pathological conditions, Tau dissociates from the microtubules and aggregates into Tau fibrils, a process ultimately leading to cellular death. However, the exact underlying molecular mechanism of Tau aggregation is still unknown.

The protein quality control (PQC) system plays a crucial role in protein folding, prevention and aggregation and controlling protein degradation. Members of the two major ATP-dependent chaperone families, Hsp70 and Hsp90, are key players in PQC. Other conserved members of the metazoan PQC network are the small heat shock proteins and the Hsp60 chaperonins, which are associated with neurodegenerative diseases (Meriin and Sherman, 2005; Webster et al., 2019).

The Hsp70-Hsp90 folding cascade has a key role in PQC and is present in various cellular compartments. The ATP-dependent chaperone Hsp70 acts as an unfoldase in the early phase of the folding cascade and after repetitive cycles of binding and release transfers its substrate for further maturation to Hsp90 (Morán Luengo et al., 2019). Hsp90 is also an ATP-dependent chaperone and acts in the decision making of its substrate, directing it either along the folding or degradation pathway (Connell et al., 2001). Next to their active role in protein folding, the chaperones also play a role in protein aggregation or disaggregation processes. The Hsp70 system acts as a disaggregation machinery for several amyloidogenic proteins (Ferrari et al., 2018; Gao et al., 2015; Kirstein et al., 2017; Nachman et al., 2019). Both Hsp70 and Hsp90 are known to interact with Tau and have a role in normal Tau regulation (Dickey et al., 2007) but also in cell stress, aggregation and degradation (Dickey et al., 2007; Kundel et al., 2018; Weickert et al., 2020), implying a role for both chaperones in AD.

If folding fails, the PQC has different degradation pathways to deal with protein misfolding and aggregation. The ubiquitin proteasomal system (UPS) degrades approximately 80-90% of proteins which are mostly short-lived, denatured or damaged (Finley, 2009). Long-lived protein aggregates are sequestered for removal by the autophagic lysosomal system (Lilienbaum, 2013). The autophagy system comprises micro-autophagy, macro-autophagy and chaperone mediated autophagy, of which the latter two are linked to the Hsp70 system. One specific type of macro-autophagy – the chaperone assisted selective autophagy – selectively degrades ubiquitin-positive substrates. The substrates are targeted towards this pathway by a complex of Hsp70:BAG3, allowing the formation of an autophagosome, which will be degraded (Arndt et al., 2007). Interestingly, BAG3 increases clearance of PolyQ aggregates via the autophagy pathway (Carra et al., 2008) raising the questions whether this could also play an important role for Tau aggregates in AD pathology.

As neurons can survive for decades despite the continuous presence of Tau protein, the PQC must be in good shape under normal conditions, in particular the Hsp70 and Hsp90 machines. This raises the question why after so many years Tau enters into a fatal aggregation process, and to which extent derailment in one or more of the PQC pathways may take centre stage in this. Multiple studies in cell lines or animal models have shown how individual components or pathways are associated with AD (Hegde et al., 2019; Klaips et al., 2018; Schaler and Myeku, 2018). Several wide-scale proteomic studies have been performed to test whether networks or pathways have altered in aging or AD (Donovan et al., 2012; Johnson et al., 2020; Sultana et al., 2007; Walther et al., 2015). It is unclear, however, whether and to which extent the PQC capacity is reduced in the human AD brain.

Recently, the Unwin group performed a wide-scale proteomics analysis on protein levels of nine AD brain in comparison to nine age-matched healthy controls (Xu et al., 2019). Their analysis revealed that an AD brain shows significant changes in specific signalling pathways, including the innate immune response and pathways involved in cell cycle regulation and apoptosis. They observe an association between extend of affectedness of brain regions and protein level changes, implying a gradual change over the course of AD pathology. Notably, the study also revealed up- or down regulation of several individual PQC factors, such as an increase in heat-shock inducible HSP70, a downregulation of the J-protein DNAJC6 and a strong increase for nucleotide-exchange factor BAG3 and small heat shock protein HSPB1. Chaperones, however, do not act as lone players, they cooperate as part of a large network. The network nature means that the PQC system could be derailed by upregulation of some factors while other members may be downregulated. Therefore, we now aim to reveal a comprehensive picture on alterations in levels of the PQC system by performing a meta-analysis on the extensive dataset provided by the Unwin group in the freely accessible Alzheimer’s Disease Proteome database (Xu et al., 2019).

We analysed a plethora of proteins who all have distinct roles in the PQC and looked for changes in any of the PQC pathways. We noticed a decrease all Hsp90 paralogues in the cytoplasm, mitochondria and endoplasmic reticulum (ER), as well as the strong upregulation of the stress-regulated pathway preparing proteins for autophagy-mediated removal. These differences are indicative of cellular distress and point towards recruitment of multiple degradation pathways for the cell trying to remove protein aggregates.

## RESULTS

### Rational of the approach

To test the hypothesis that the PQC capacity decreases in AD, we performed a meta-analysis of the proteomics data provided by the Unwin laboratory, which is freely available at http://www.dementia-proteomes-project.manchester.ac.uk/Proteome/Search (Xu et al., 2019) (Fig. 1). This study reveals a quantitative overview on protein levels of nine AD brains and nine healthy age-matched control brains, separated per brain region. The six distinct brain regions studied range from mostly unaffected in AD (cerebellum) to mildly affected (motor cortex and sensory cortex) and strongly affected (hippocampus, entorhinal cortex and cingulate gyrus) (Smith, 2002; Xu et al., 2019). We analysed 94 distinct proteins involved in different pathways of the PQC to obtain a comprehensive overview of protein levels of components of the PQC in AD (Fig. 1). AD levels are represented as fold-change compared to control brain. Significance of increase or decrease in protein levels is determined by assessing the false-discovery rate (FDR), indicating significant false positive values. Generally, FDR values provide an estimate on variability of the data. An FDR of e.g. 1% indicate 1 incorrect positive hit per 100 values with an FDR below 1%.

**Figure 1.**
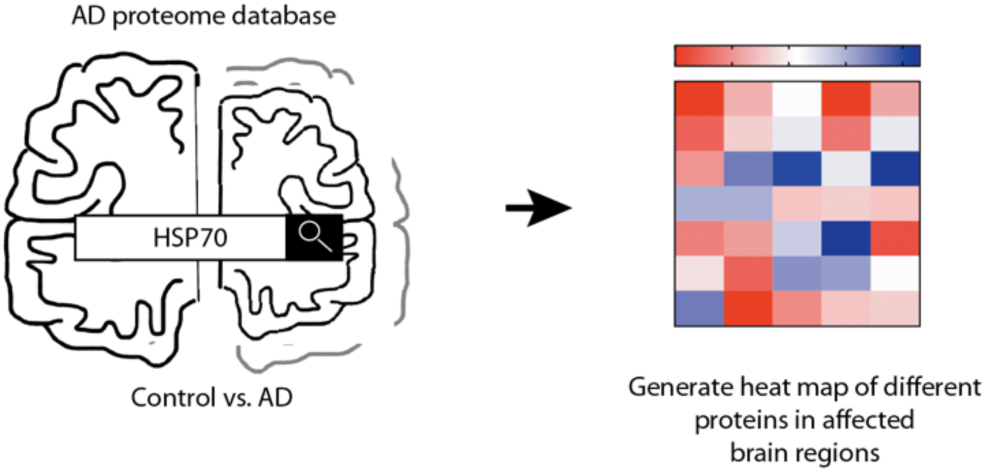
Schematic representation of approach. Schematic overview of rational of our analysis. Protein expression levels were obtained from the Alzheimer’s Disease Proteome Database and compared between 9 AD brains and 9 age-matched control brain. Heat map was created to illustrate the protein level changes according to brain region.

### Neuronal and glial cells

Brain tissue consists of multiple cell types, which either are neuronal cells or glial cells. While neuronal degradation is at the heart of the disease, also glial cells are implied to have a role in AD (Dzamba et al., 2016). To assess whether the database allows conclusions on the PQC system in all different cell types, we analysed for the presence of cell-type specific protein markers (Fig. 2). For glial cells at least one specific protein markers is present for each of the three types of glial cells; Iba1 for microglia, GFAP for astrocytes and MPB for oligodendrocytes (Fig. 2A), indicating that all of them are still present in AD brain (Imai et al., 1996; Nawaz et al., 2013; Pekny and Pekna, 2004). Even though none of the protein levels are below the 1% FDR threshold value, for microglia and astrocytes we do observe a pattern related to disease progression with relatively small FDR value. Microglia marker levels are increased with 16-26% in hippocampus and entorhinal cortex, and astrocytes increase with 38%-44% in all three affected brain regions, possibly indicative of an overrepresentation of these type of cells in affected brain regions. For neurons, we also assessed the presence of markers for the brain’s most prominent excitatory and inhibitory neurons; VGLUT for glutamatergic and GAT1 for GABAergic neurons (Borden, 1996; Fremeau et al., 2001). Both proteins were present in all six brain regions (Fig. 2B), indicating that these neurons are also represented in the database.

**Figure 2.**
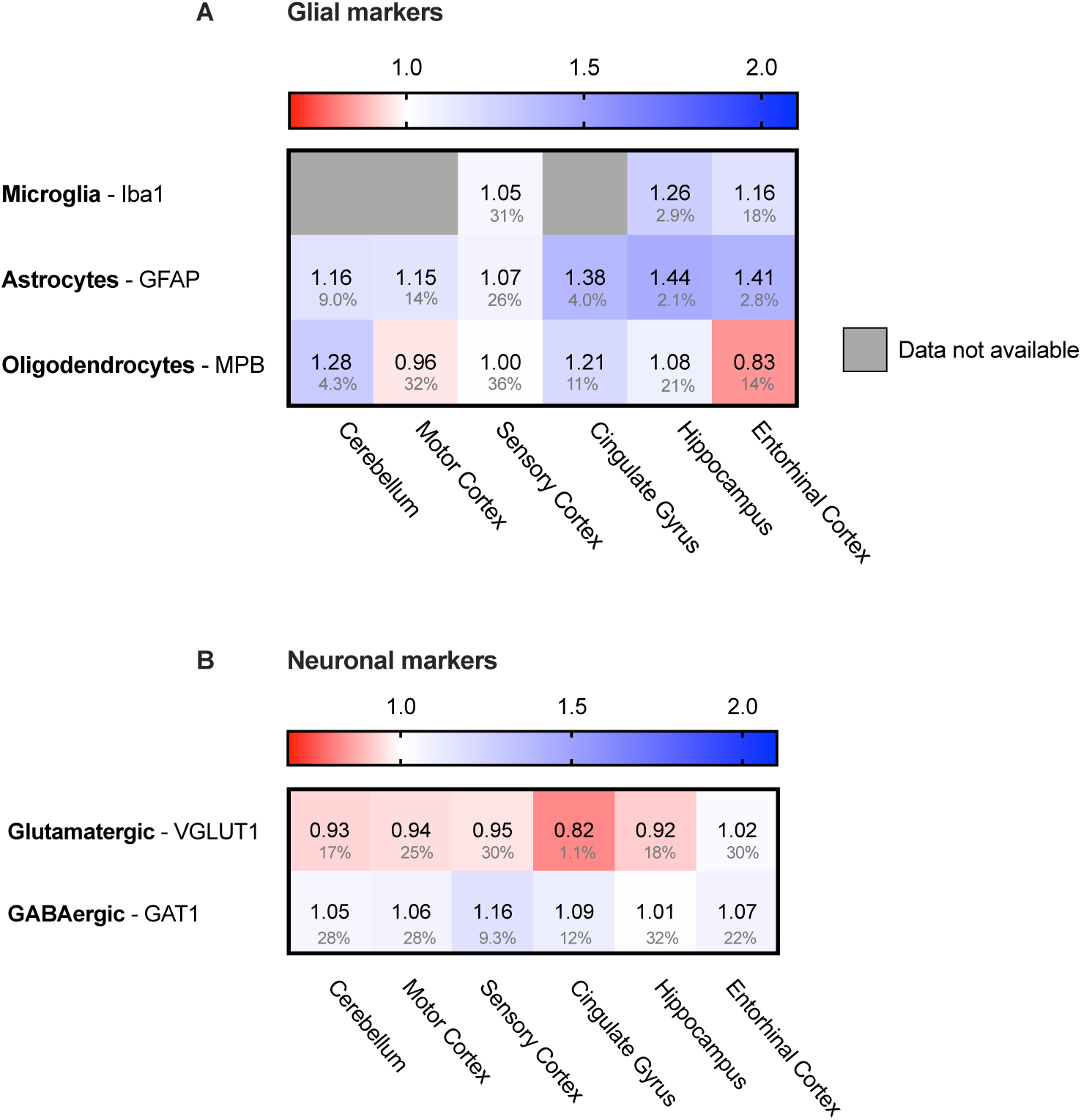
Both neuronal and glial cells present in analysed sample. Heat map of neuronal and glial protein markers in different brain regions, differently affected in Alzheimer’s Disease. Protein levels are calculated as fold change compared to control brain, with baseline set at 1.0 in white (averaged numbers from nine Alzheimer patients and nine controls of similar age, based on the proteome analysis of Xu et al (Xu et al., 2019) increase in protein levels, blue gradient; decrease, red gradient). Grey boxes represent unavailable data. Actual numbers of fold change are indicated in corresponding boxes, FDR values are indicated as percentage below. **A**. Protein markers for all glial cells are present in all six brain regions, indicating presence of these types of cells in the analysed sample. **B**. protein markers for glutamatergic (excitatory) neurons and GABAergic (inhibitory) neurons, indicating presence of both type of neurons in analysed brain tissue.

### Levels of aggregating Tau are not affected in AD

As Tau fibrils are a major hallmark of Alzheimer’s Disease, we wondered whether the appearance of tangles may be reflected in potentially higher levels of Tau in AD brains. Analysis of Tau protein levels between AD brains and control, however, showed no major differences in protein levels in any of the brain regions (Fig. 3). This indicates that Tau fibril formation is not driven by an increase total Tau levels.

**Figure 3.**
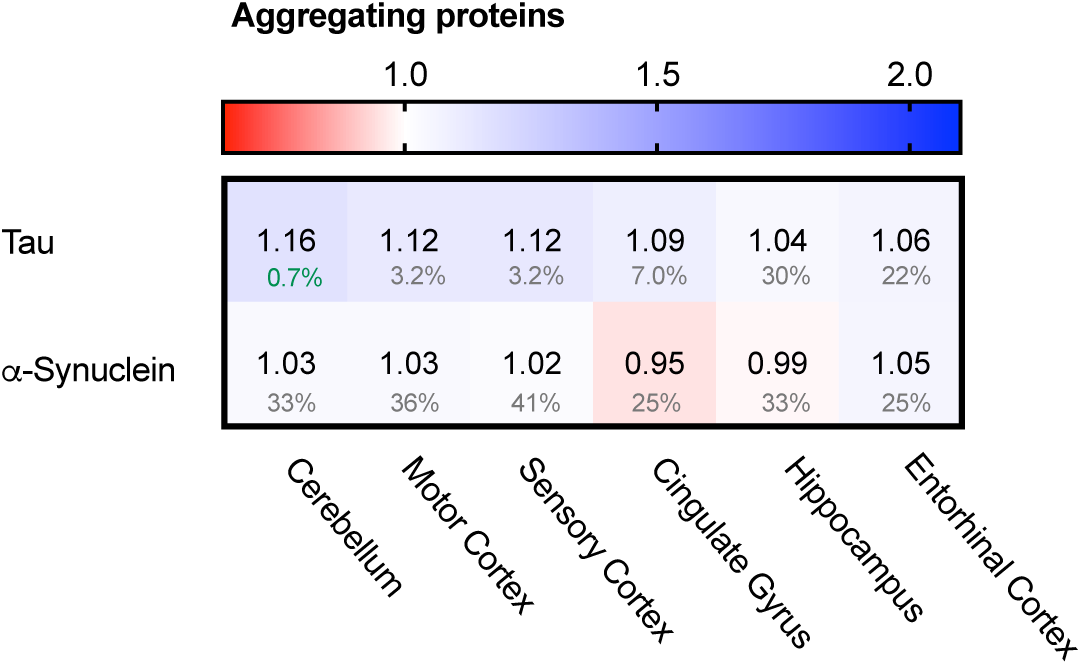
Levels of Tau and α-synuclein not affected in Alzheimer’s Brain. Heat map of protein levels in different brain regions, differently affected in Alzheimer’s Disease; colour code as in Fig. 2. Tau levels remain largely unaffected with only minor increase in some brain regions, whereas α-synuclein remains unaffected in all brain regions.

We also looked into the levels of a-synuclein. Alpha-synuclein is an aggregating protein in Parkinson’s Disease and Fronto-Temporal Dementia but is also involved in AD via crosstalk with Tau in promoting each other’s aggregation (Attems and Walker, 2017). When analysing protein levels of a-synuclein we could not identify notable differences in the distinct brain regions (Fig 3). Together, there are no noteworthy differences in the levels of two major proteins that aggregate in disease. This suggests that the key difference in the AD brain is not related to changes in the levels of the aggregating proteins themselves. Therefore, we set out to investigate whether disturbance of the PQC network, which controls and prevent protein aggregation in healthy neurons, may be a hallmark of neurons in AD.

### Chaperonins do not alter in AD

We started the analysis of the PQC system with the largest folding machine in the metazoan cytosol, the ATP-dependent HSP60 chaperone family, also known as chaperonins (Ansari and Mande, 2018). The chaperonin TRiC/CCT is associated with protein aggregation in disease, in particular in Huntington’s Disease, where it can bind to specific subunits of the huntingtin protein and modulate its aggregation (Spiess et al., 2006; Tam et al., 2009). We wondered whether chaperonin levels are affected in AD and analysed the levels of the seven TRiC/CCT subunits in the cytoplasm and HSP60 in mitochondria. None of the TRiC/CCT subunits showed notably altered levels in the Alzheimer brain, ranging from only 10% for CCT2 in the entorhinal cortex to no change at all in the motor cortex for CCT7 (Fig. 4). Similarly, HSP60 levels did not differ between Alzheimer and control brains, as values change from a minor decrease of 1% in the hippocampus to a slight 6% decrease in the cerebellum (Fig. 4).

**Figure 4.**
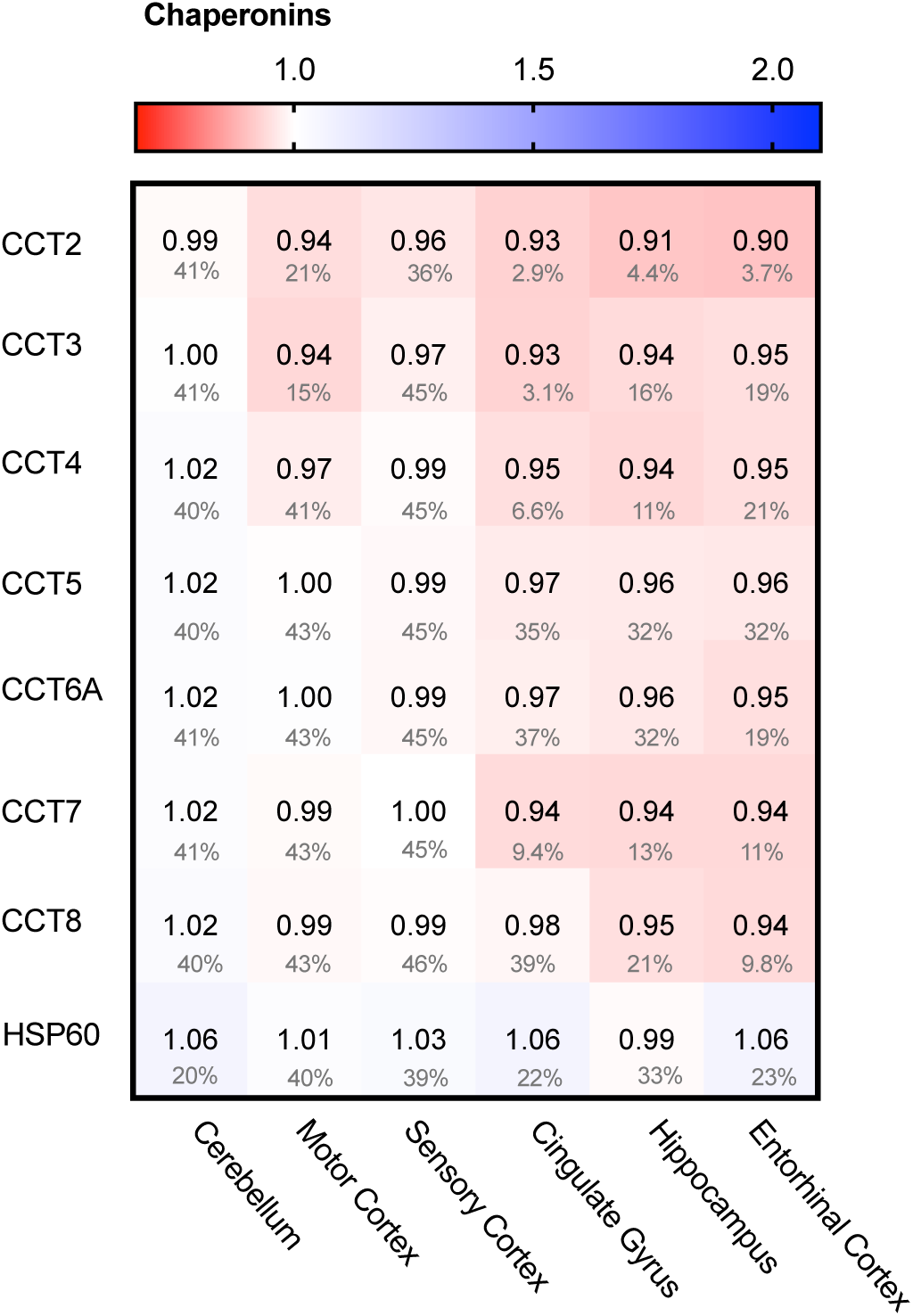
Chaperonin levels do not alter in AD. Heat map of chaperonins in different brain regions; colour code as in Fig. 2. None of the TriC/CCT subunits show notable differences in protein levels. Mitochondrial HSP60 levels also remain fairly constant in all brain regions in AD.

Alterations in chaperonin levels thus do not to play an important role in AD.

### Decrease of all Hsp90 paralogues in AD

One of the major chaperone systems controlling Tau is the evolutionary conserved Hsp90 machinery. To see if Hsp90 levels are affected in AD brain, we compared in the database the levels of all four Hsp90 paralogs. Strikingly, all Hsp90 paralogs were decreased in all brain regions affected in AD (Fig. 5A).

**Figure 5.**
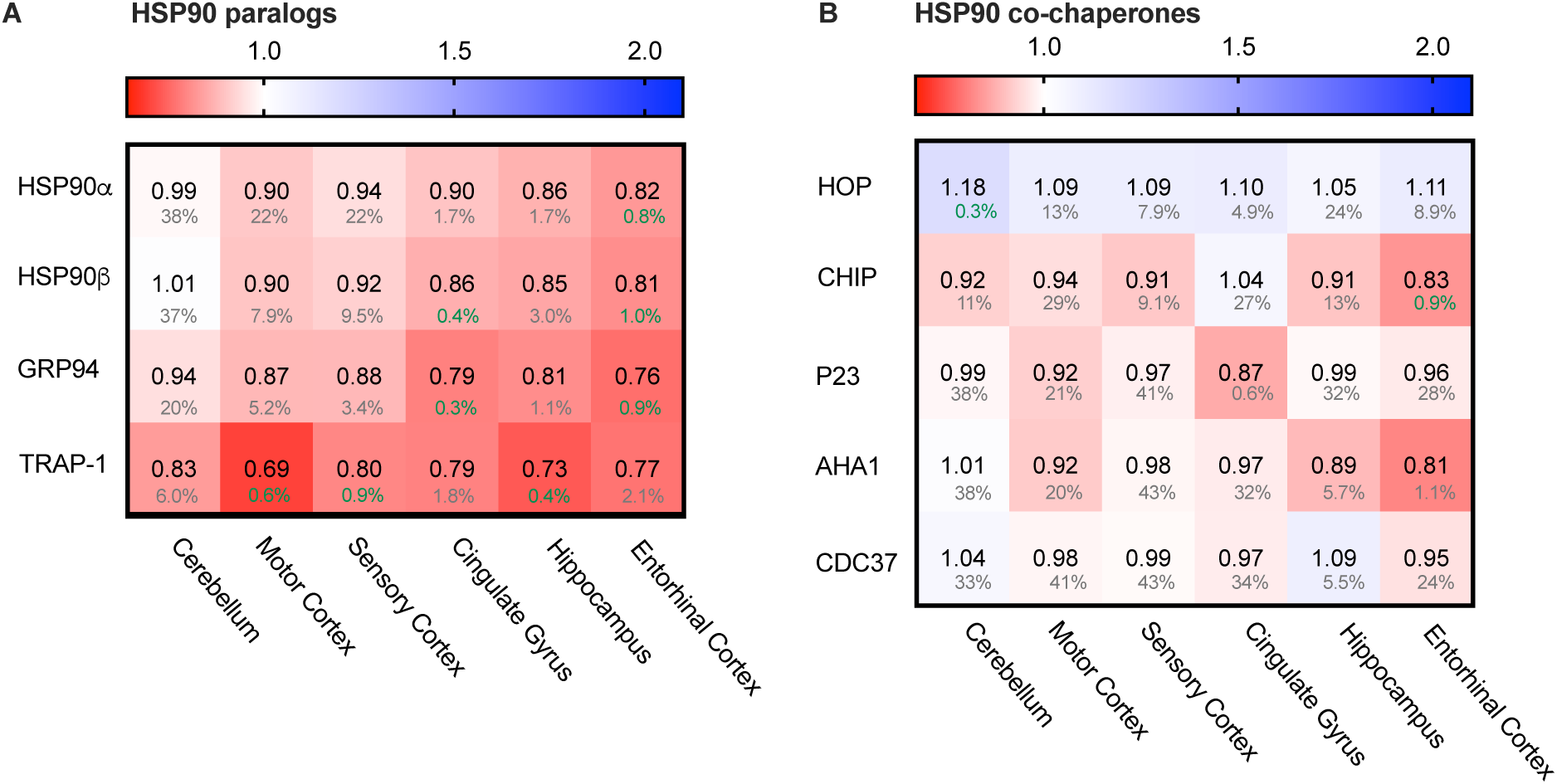
All HSP90 paralogues decreased in AD. Heat map of HSP90 paralogues and it’s co-chaperones in different brain regions; colour code as in Fig. 2. **A.** All HSP90 paralogs show a decrease in protein levels, with the strongest decrease in mitochondrial HSP90, TRAP-1. The cerebellum, which is the most unaffected brain region in AD, appears to be mostly unchanged, except for TRAP-1 levels. **B.** Co-chaperones of HSP90 show no strong differences between control and AD brain.

Of all Hsp90s, the mitochondrial TRAP-1 showed the most pronounced decrease in Alzheimer neurons, with a drastic decrease of 31% in the motor cortex. The other AD affected brain regions also showed severe reduction in TRAP-1 levels, such as a decrease of 27% in the hippocampus and 21% in cingulate gyrus. For the cytosolic house-keeping paralogue HSP90b, levels decreased with 18% in the entorhinal cortex in AD compared to control, and heat-shock inducible HSP90a showed a similar reduction in the same brain region (19%). Interestingly, for all paralogues the levels in the cerebellum do not show notable changes in AD. The cerebellum is the brain region least affected in AD, which makes it likely that the decrease of Hsp90 levels in the more disease-affected brain regions is Alzheimer-related.

To place the relative reduction levels into context, it is important to realise that the changes documented in the database represent the averaged protein levels in the entire tissue. Some cells may be severely depleted while others remain fairly unaffected. This makes it likely that actual changes in Hsp90 levels in AD-affected cells are more drastic than reflected in the percentage values in the database.

The function of cytoplasmatic Hsp90 is specified by a plethora of co-chaperones. We looked, therefore, also into protein levels of several co-chaperones (Fig. 5B). In contrast to the Hsp90s, all differences in co-chaperone levels were only minor (4% for p23 in entorhinal cortex and 8% for Aha1 in motor cortex) and were accompanied by high FDR rates, which render them insignificant. Thus, although all Hsp90s itself is depleted in AD, its regulatory network is not derailed.

### Only heat shock inducible HSP70 strongly enhanced in AD

Hsp90 depletion is known to upregulate HSF1 (Zou et al., 1998) which is considered to be the master regulator of the heat shock response and is involved in the regulation of Hsp70 expression. Next to Hsp90, Hsp70 is the other conserved ATP-dependent chaperone family that is present in all folding compartments. Hsp70 acts upstream of Hsp90 in the early stage of the folding cascade and is known to interact with Tau (Dickey et al., 2007). We evaluated the levels of four Hsp70 paralogues: HSP70 and HSC70 in the cytoplasm, mt-HSP70 in mitochondria and BiP in the endoplasmic reticulum. All these paralogs are constitutively expressed, except for the cytosolic heat-shock inducible HSP70, which is under the regulation of HSF1. Remarkably, heat-shock inducible HSP70 was strongly increased in AD brain of all brain regions affected in disease (e.g. 24% in the hippocampus and entorhinal cortex and 18% in cingulate gyrus), whereas the constitutively expressed paralogues remained constant (Fig. 6A). As HSP70 is under regulation of HSF1, it is involved in the stress-response of the cell and its upregulation may indicate derailment of the cellular stress response.

**Figure 6.**
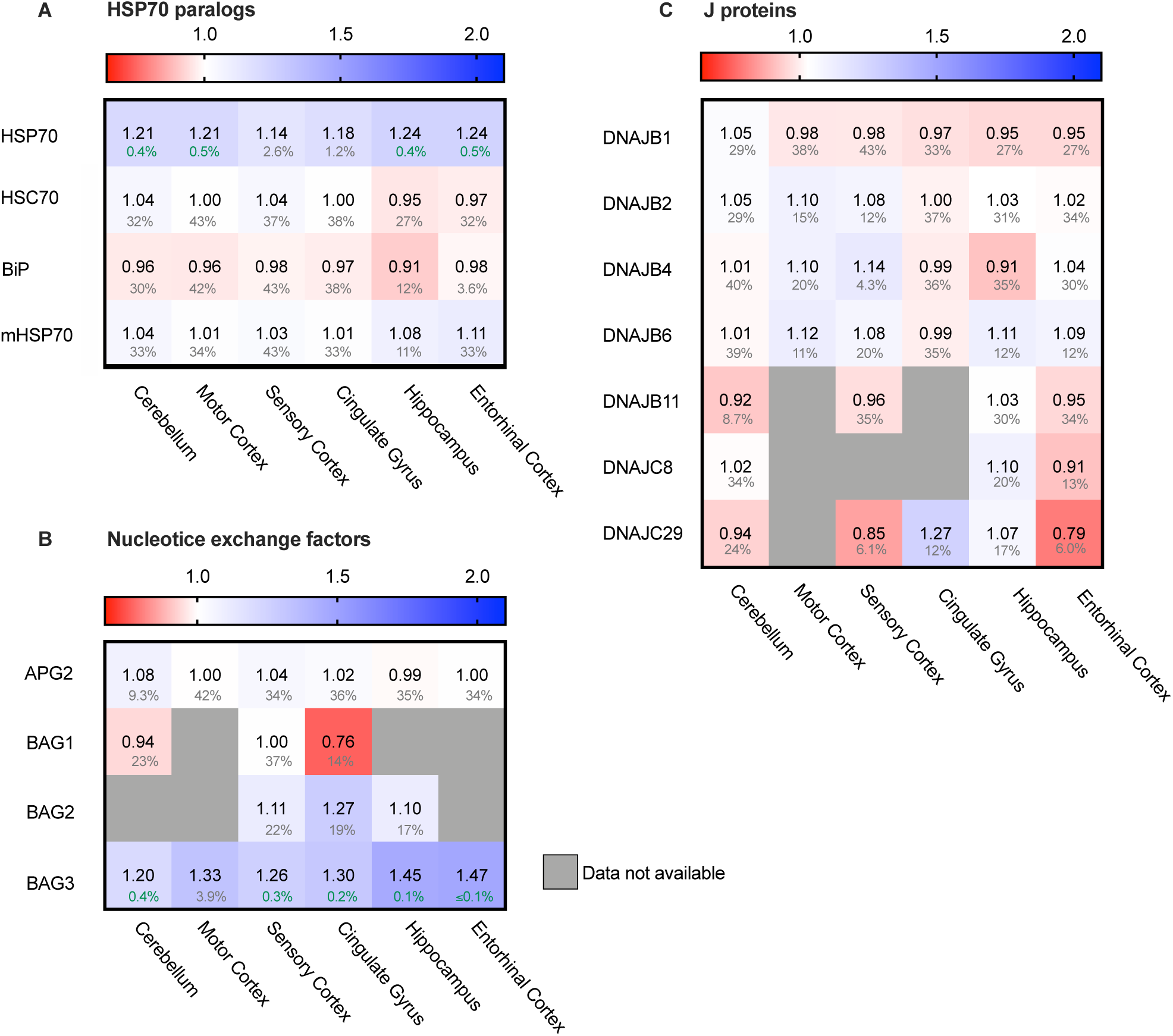
Only heat-shock inducible HSP70 strongly increased in AD. Heat map of paralogs of HSP70 and its co-chaperones in different brain regions; colour code as in Fig. 2. **A.** Heat shock inducible HSP70 is strongly upregulated in all brain regions, whereas all other paralogs of HSP70 remain constant. **B.** J-proteins do not show a general tendency in increase or decrease. DNAJC29 is decreased in sensory cortex and entorhinal cortex, but a high false discovery rate renders these changes not significant. **C.** Analysed NEFs of HSP70 are unaffected, except for BAG3. BAG3 is strongly increased in all brain regions.

### HSP70 efflux derails towards autophagy

Hsp70s are functionally dependent on co-cochaperones. We looked into a subset representing the most prominent J-proteins and Nucleotide Exchange Factors. J-proteins stimulate the Hsp70 ATPase and act as substrate targeting factors for Hsp70s, thereby controlling the influx into this chaperone system. We evaluated the protein levels of J-proteins associated with neurodegenerative diseases (Kampinga and Craig, 2010). Notably, we did not observe major differences for any of the J-proteins (Fig. 6C). The most notable change was an increase of 14% for DNAJB4 in the sensory cortex, but this co-chaperone did not show a general pattern in relation with AD affected brain regions. Overall, the levels of J-proteins did not show prominent differences between Alzheimer and control neurons, which makes it unlikely that control of substrate influx into the Hsp70 system is disturbed in AD.

After ATP hydrolysis of Hsp70, the subsequent replacement of ADP by ATP releases the substrate protein. Nucleotide Exchange Factors (NEFs) trigger this exchange, thereby controlling environment and conditions of release and resetting the Hsp70 system (Mayer and Gierasch, 2019). Thus, NEFs regulate substrate efflux, triaging the fate of the substrate after its release from Hsp70, including refolding, disaggregation and degradation either by the proteasome or by autophagy. A key NEF for folding and disaggregation is APG2 (Bracher and Verghese, 2015). It contributes to the disaggregation capacity of Hsp70 for aggregates of Tau, a-synuclein and huntingtin (Ferrari et al., 2018; Gao et al., 2015; Kirstein et al., 2017). In AD brains however, its levels remained unaffected (Fig. 6B). Strikingly, levels of another NEF, BAG3, are strongly elevated in AD brain (Fig. 6B). Interestingly, BAG3 has a specific role in chaperone assisted selective autophagy by forming a multi-chaperone complex for ubiquitylation and sequestration of its client protein and facilitates in the substrate engulfment by the autophagosome (Klimek et al., 2017). Its expression is part of the cellular stress response and controlled by HSF-1 (Franceschelli et al., 2008). Levels of its family members BAG1 and BAG2 were not decisively altered (Fig. 6B). The strong upregulation of BAG3 in AD brains, in striking contrast to its family members, implies a change in the efflux control of the Hsp70 machinery towards activation of the autophagy pathway.

### Upregulation of several sHSPs

In contrast to the ATP-driven Hsp70 and Hsp90 systems, small heat shock proteins (sHSPs) are the largest ATP-independent class of chaperones. They act early-stage on hydrophobic stretches of their substrates, possibly upstream of the ATP-dependent Hsp70-Hsp90 chaperone cascade (Bakthisaran et al., 2015; Haslbeck and Vierling, 2015; McHaourab et al., 2009; Morán Luengo et al., 2019; Zwirowski et al., 2017). Out of 10 mammalian sHSPs, five family members are expressed in neurons and play a role in neurodegenerative diseases: HSPB1, HSPB5, HSPB8 and, to a lower extend, HSPB6 (Quraishe et al., 2008; Webster et al., 2019). HSPB1 and HSPB6 both show a strong increase in all brain regions (Fig. 7). Similar trends can be observed for HSPB5 and HSPB8, although they are accompanied by slightly higher FDR values and may thus not be fully representative. HSPB1 levels increased from only 5% in the unaffected cerebellum up to 46% in the hippocampus. For HSPB6 levels increased even further up to 66% in the cingulate gyrus. sHSPs co-operate with HSP70 and BAG3, the two other chaperone components also strongly increased in AD brain (Fig. 5, (Rauch et al., 2017)). The sHSP-HSP70-BAG3 system channels its substrate towards chaperone assisted selective autophagy (Stürner and Behl, 2017). Thus, within the entire cellular chaperone network only the pathway leading towards autophagy is upregulated in AD.

**Figure 7.**
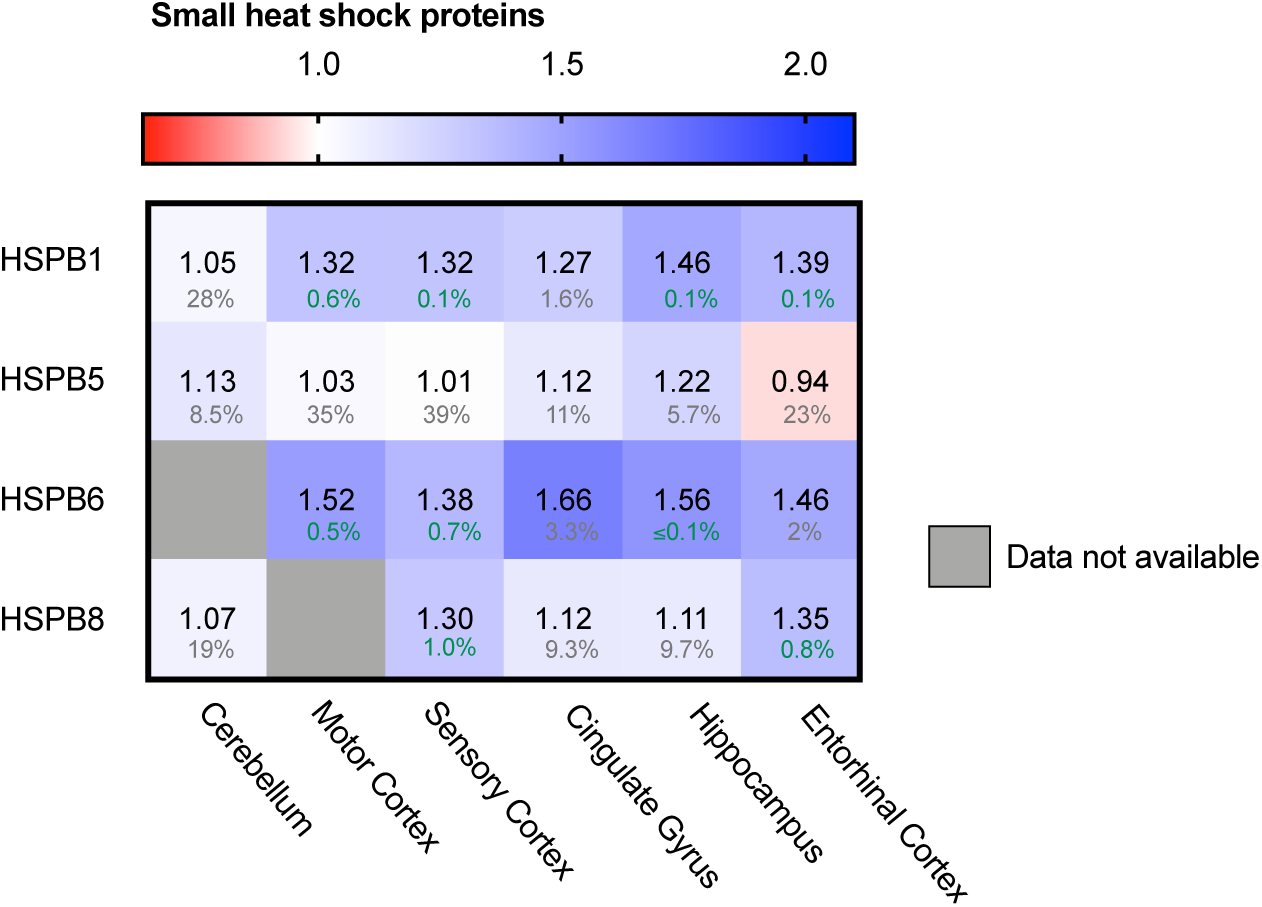
Neuronally expressed sHSPs upregulated in AD brain. Heat map of sHSPs in different brain regions; colour code as in Fig. 2. HSPB1 and HSPB6 are strongly increased in all brain regions, HSPB5 and HSPB8 show only minor differences accompanied by high FDRs.

### Alterations in autophagy markers in Alzheimer brains

Autophagy is the degradation pathway dedicated to protein aggregates and dysfunctional cellular compartments (Finley, 2009). In particular, selective macro-autophagy targets such aberrant components, engulf them in double membrane compartments termed autophagosomes. Subsequent fusion with lysosomes allows enzymatic degradation (Stolz et al., 2014). Protein aggregates can be cleared by selective autophagy - a process referred to as aggrephagy – and could possibly be of relevance in AD. LC3 proteins are early-stage markers for autophagy and are important for the autophagosome formation. Map1LC3A levels were decreased in AD brain (Fig. 8A, 16% in entorhinal cortex), indicating impairment in this process. Several adaptor proteins are known for autophagy, with different adaptor proteins specific for individual autophagy pathways. Adaptor protein SQSTM-1 is key marker for aggrephagy (Zaffagnini et al., 2018) and showed an extreme increase in protein levels in the brain region most affected in AD; the entorhinal cortex (111%, Fig. 8B). Increased levels of SQSTM-1 are indicative of impaired autophagy (Bjorkoy et al., 2005) implying that this process is impaired in AD. More specifically, increase of SQSTM-1 is likely indicative of recruitment of SQSTM-1 mediated aggrephagy in AD, as other autophagy adaptors, such as OPTN, are not meaningfully altered in AD.

**Figure 8.**
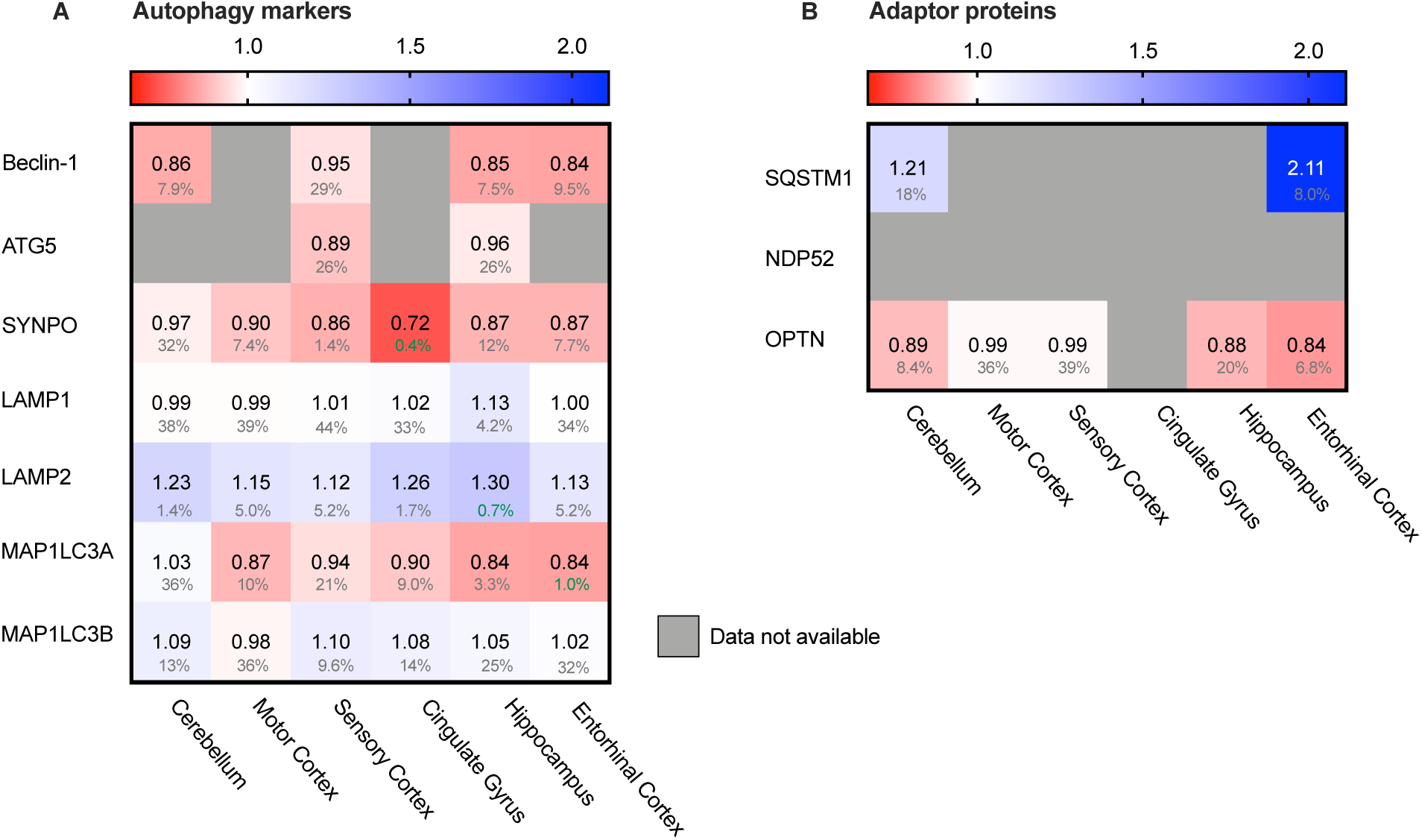
Chaperone-mediated autophagy adaptor SQSTM1 increased in AD. Heat map of autophagy markers and their adaptor proteins in different brain regions, colour code as in Figs. 1 and 2. **A.** General autophagy-markers do not show any significant difference, except for MAP1LC3A in the strongest affected brain regions (Hippocampus, Entorhinal Cortex and Cingulate Gyrus). Observed differences in other brain regions are non-representative due to high false discovery rate. **B.** The protein SQSTM1, adaptor for autophagy, is increased in both brain regions for which data are available.

### Proteasomal degradation system remains largely unaffected

The main cytosolic degradation pathway for many processes is the ubiquitin-proteasomal machinery, including for the Alzheimer protein Tau. A cascade of E1, E2 and E3 ligases target proteins for degradation by flagging them with a poly-ubiquitin chain (Zheng and Shabek, 2017). When analysing levels of several different E1, E2 and E3 ubiquitin ligases in the Alzheimer brain, we did not note significant differences. For small subset of ubiquitin ligases we did observe minor decreases, but these were all accompanied by high FDR values, rendering them insignificant (Fig. 9A-C). E1 ubiquitin ligase UBA1 for instance was decreased by 22% in entorhinal cortex and SAE even by 30% in the motor cortex (Fig. 9A). However, high FDR values mark them as likely outliers.

**Figure 9.**
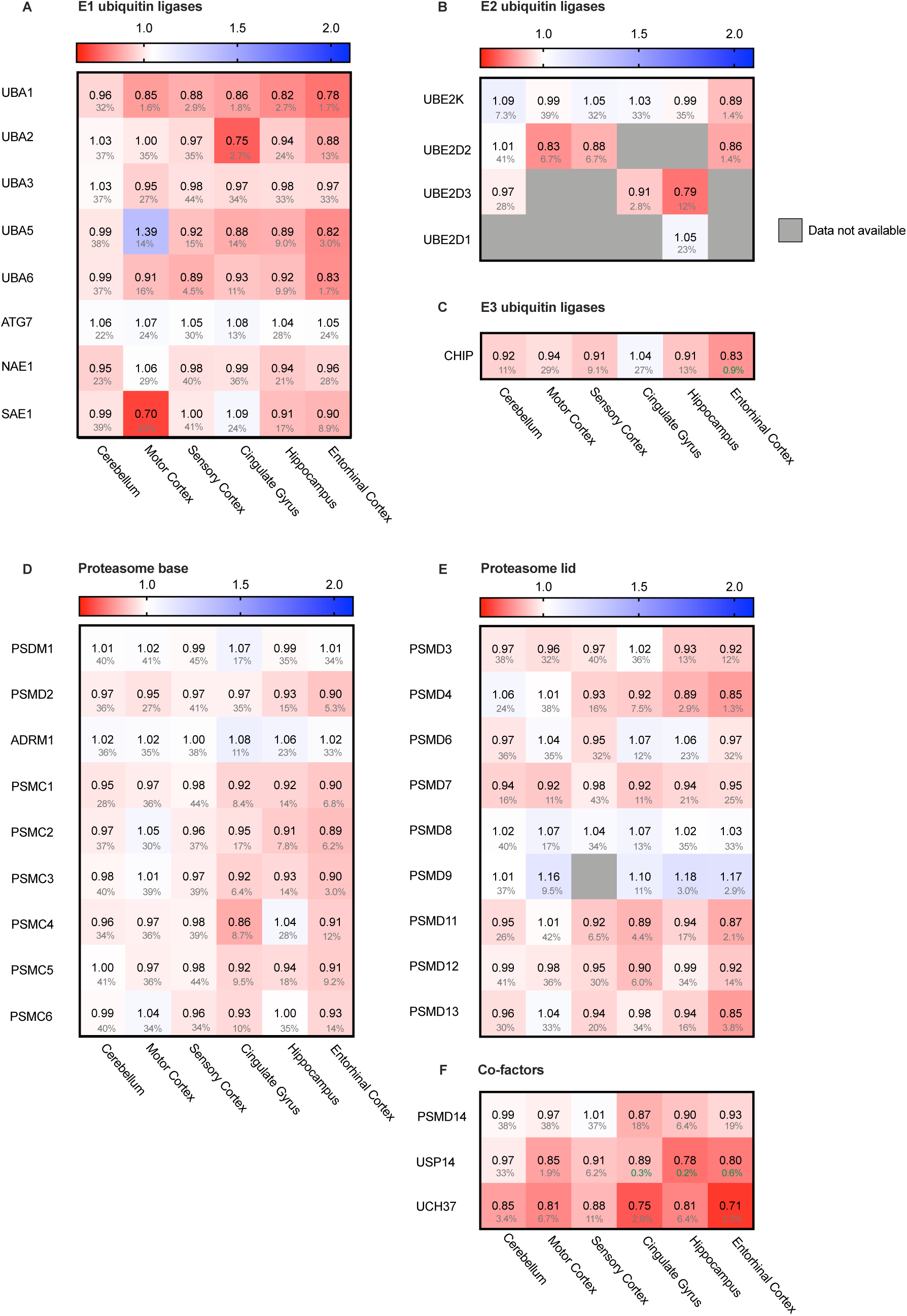
Ubiquitin ligases largely unaffected in AD. Heat map of components of the ubiquitin ligases in different brain regions; colour code as in Fig. 2. **A-C**. E1, E2 and E3 ubiquitin ligases do not show any notable difference in protein levels in AD affected brain regions. **D-E**. Components of the base and lid of the proteasome do not show any notable difference in protein levels in any of the brain regions. **F.** Some co-factors of the proteasome, such as Usp14 and Uch37, are slightly decreased in AD brain.

Next, we turned from the targeting cascade to the degradation machinery itself and analysed the different subunits of the proteasome. The proteasome is build-up of two complexes: a four-ring core component and one or two regulatory lid structures (Coux et al., 1996). Neither for the proteins constituting lid nor base of the proteasome we did noted differences in protein levels (Fig. 9D,E). Interestingly, however, two of the three metazoan de-ubiquitinating enzymes associated with the proteasome (de Poot, S A H et al., 2017), USP14 and UCH37, showed decreased levels throughout all brain regions, ranging from 13-29% (Fig. 9F). These factors trim the ubiquitinated substrate on the proteasome, slowing down proteasomal degradation (Lam et al., 1997). Inhibition of these factors upregulates proteasomal activity (Lee et al., 2011). Decreased levels of these co-factors in AD brains could thus indicate increase of proteasomal flux.

### Hsp90 only stress system affected in mitochondria

Protein folding is a process not restricted to the cytosol, but also occurs in other cellular compartments such as the mitochondria. Mitochondria are essential for energy supply, but also play a key role in activation of cellular apoptosis and thus cellular degeneration. We therefore analysed four mitochondrial chaperones (Fig. 10A). HSP60, CLPB and mt-HSP70 did not show major differences in protein levels. The Hsp90 paralog TRAP-1 is thus the only mitochondrial chaperone strongly decreased in all brain regions. This downregulation of TRAP-1 reflects a specific pathway, as a general stress response would most likely affect levels of other chaperones in mitochondria as well.

**Figure 10.**
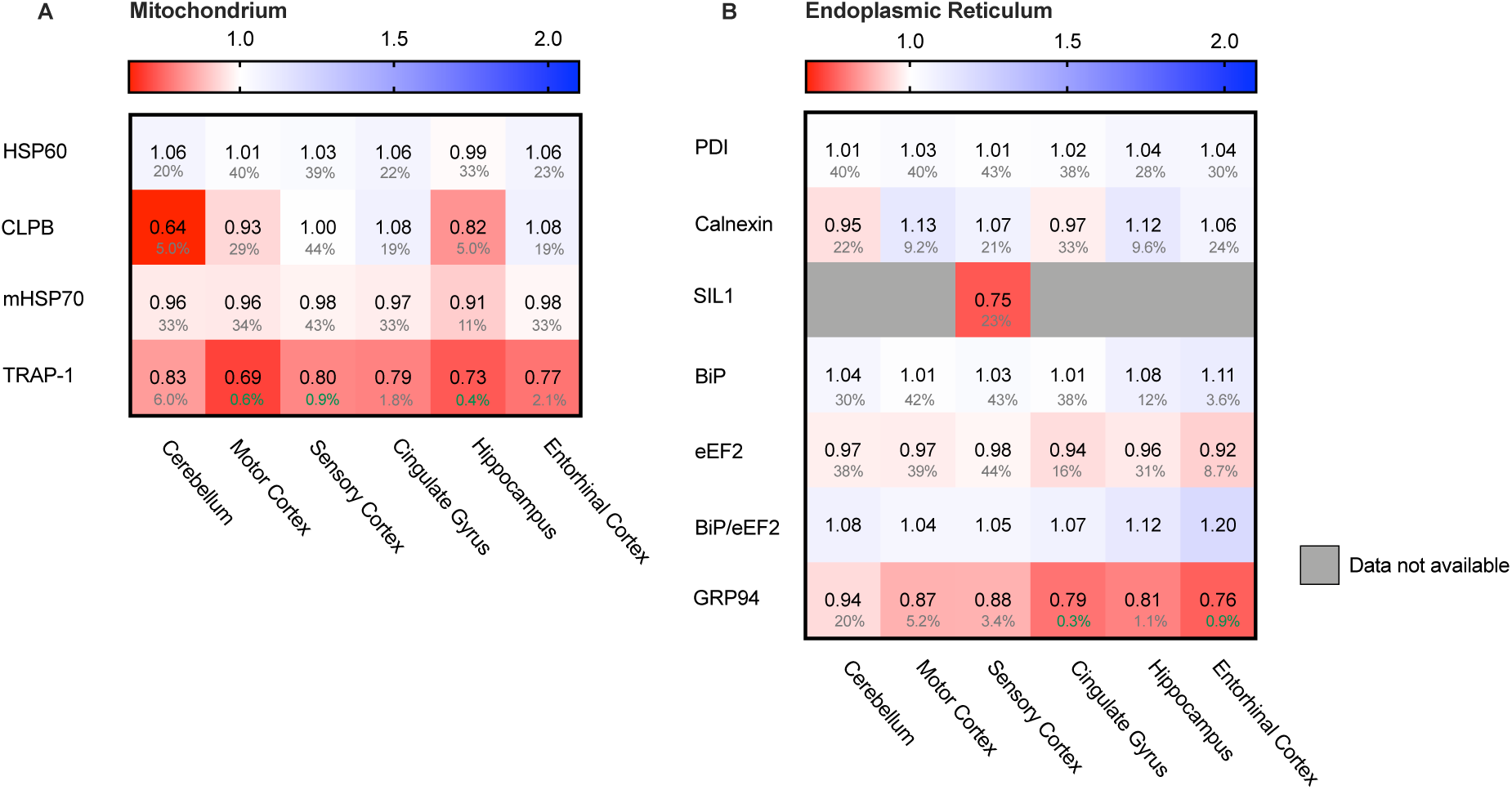
HSP90 paralogue TRAP-1 strongly decreased in AD. Heat map of mitochondrial and ER chaperones in different brain regions; colour code as in Fig. 2. **A.** The HSP90 paralogue, TRAP-1, shows a strong decrease in protein levels in AD. Other mitochondrial chaperones, including mitochondrial HSP70, HSP60 or CLPB, do not show any notable changes in protein levels in AD. **B.** Strong decrease in protein levels of GRP94, the HSP90 ER paralogue. Other components remain fairly unaffected. The decrease in SIL1 protein level most likely is not representative because of high false discovery date.

### Endoplasmic Reticulum chaperones does not show remarkable alterations

The endoplasmic reticulum (ER) is a cellular compartment important in protein production and folding. ER stress leads to the activation of the Unfolded Protein Response (UPR), implied to be upregulated in AD in the hippocampus and entorhinal cortex (Cornejo and Hetz, 2013; Hoozemans et al., 2005). The ER paralog of Hsp90 - Grp94 - does show a strong decrease, ranging from a minor 6% in the mostly spared cerebellum, up to an astonishing 23% in the highly affected entorhinal cortex (Fig. 10B). Individual levels of the main luminal Hsp70, BiP, shows no major changes. The levels of the Hsp110 SILl-1 NEF for BiP, are also not affected in any of the brain regions, with the exception of a 25% decrease in the sensory cortex. However, this is not representative, as this finding is restricted to only one brain region and accompanied by a high FDR of 23%. Other UPR markers, such as PDI and Calnexin, show no major differences in any of the brain regions in AD. As there is no general increase of UPR specific markers we conclude that the Alzheimer brain is not characterised by a fully activated UPR.

## DISCUSSION

In this paper we aimed to reveal systematic changes in the PQC network in Alzheimer neurons (Fig. 11A). Interestingly, the majority of players in the field remain unaffected, but a subset of the PQC components is either up- or downregulated in AD. Levels of all Hsp90 paralogues in cytoplasm, mitochondria and ER are strongly decreased. In contrast, a chaperone system consisting of heat shock inducible HSP70, its NEF BAG3 and several sHPSs shows remarkable increases in protein levels (Fig. 11A).

**Figure 11.**
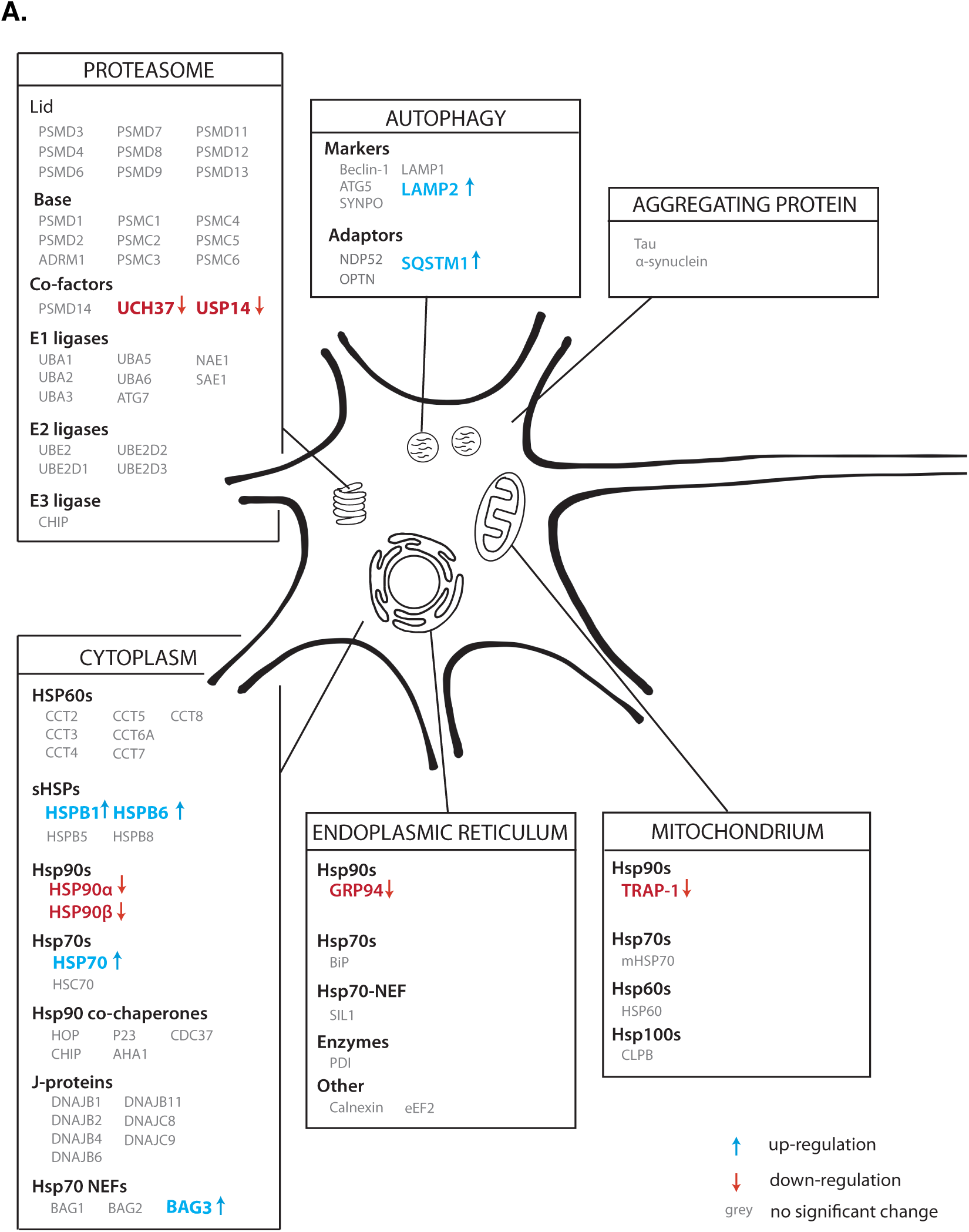
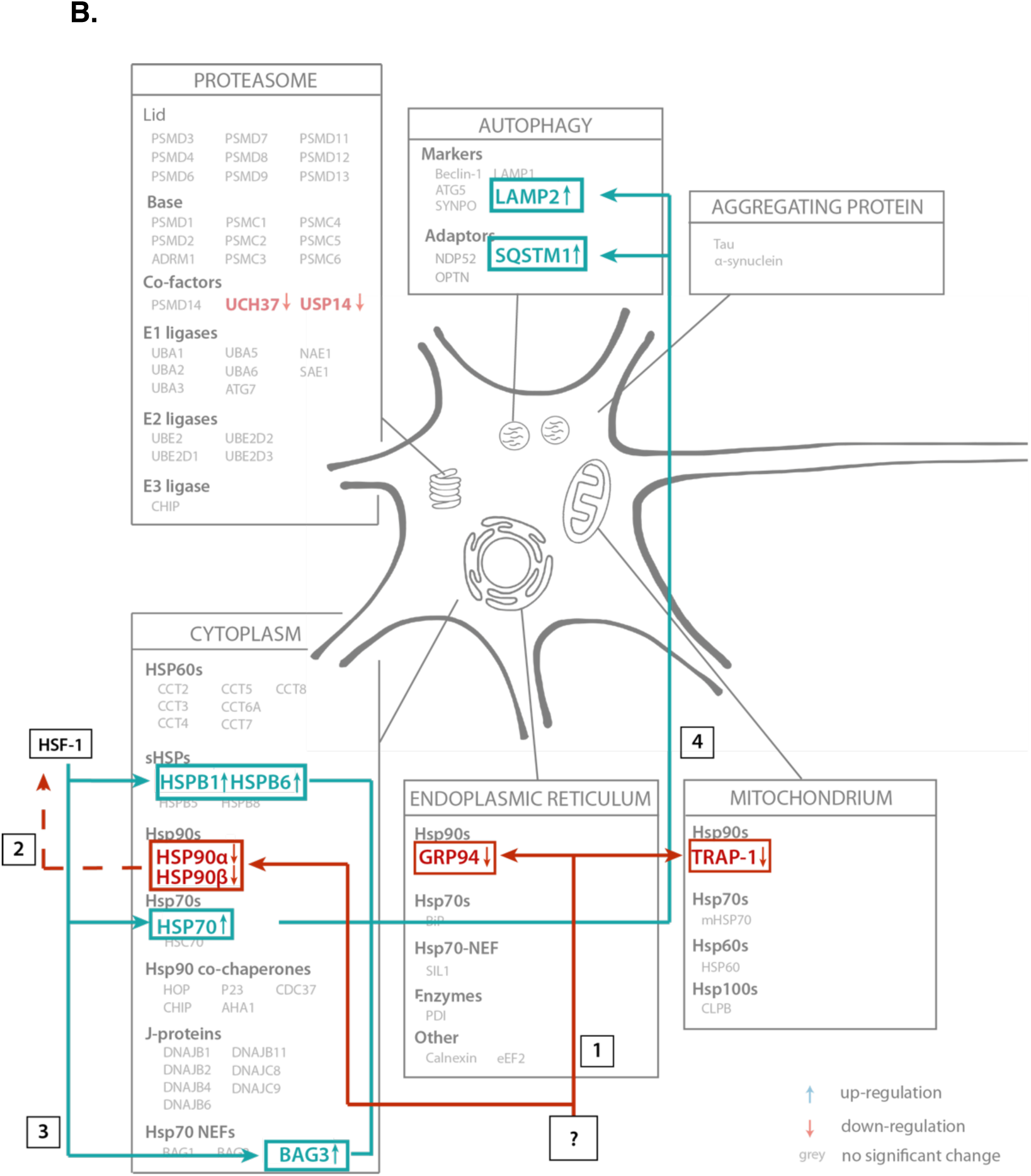
Illustrative overview of main findings. **A.** Schematic representation of the alteration of protein levels in our analysis. Proteins are sorted per compartment and classification. Proteins represented in grey are not significantly altered in AD brains, proteins in blue are upregulated and proteins in red are downregulated. **B.** Functional connection of PQC factors affected in AD. An unknown event ‘?’ causes downregulation of all HSP90 paralogs (1), which may lead to activation of transcription of heats shock genes, possibly via HSF-1 (2), leading to subsequent upregulation of HSP70, BAG3 and sHSPs (3). These factors then trigger activation of chaperone assisted selective autophagy factors SQSTM-1 and LAMP2 (4).

### Neurons versus glial cells

Our findings are based on analysing the Alzheimer Disease Proteome database from the point of view of the PQC system (Xu et al., 2019). Thus, the accuracy of our conclusions depends on the quality of the data provided. While the data specify protein levels per brain region, they do not resolve different cell types with these regions. The brain material from which the samples are taken consists of both glial and neuronal cells, and protein markers of both types of cells were present in the analysed tissue. Therefore, both neurons and glial cells may contribute to the measured protein levels (Fig. 2). It is evident from our analysis that the deviations in the Alzheimer tissue reflect a specific adaptation of a particular cellular stress pathway, raising the question which cell type may be affected most. There are three options: (i) The stress response is exclusive for neurons; (ii) it is exclusive for glial cells; and (iii) the response takes place in both neuronal and glial cells. There is conflicting evidence whether specific sHsps are expressed more in neuronal or glial cells. (Björkdahl et al., 2008; Schwarz et al., 2010; Wilhelmus et al., 2006). HSPB1 and HSPB5 are upregulated in neurodegenerative diseases, especially in reactive glial cells and HSPB8, together with BAG3, is upregulated in astrocytes (Seidel et al., 2012). Both microglia and astrocytes are activated in the neuroinflammatory response and implied to contribute to AD pathology (Kaur et al., 2019; Stürner and Behl, 2017). However, as the neurons are the cells that are suffering from intracellular protein aggregation, it seems likely that the observed changes in protein quality control represent a response inside neurons as well. Given that in the cortex, which is most affected in Alzheimer, glial cells outnumber neurons by 4:1, the alterations in PQC level would underestimate the reaction at neuronal level. In contrast, the second alternative would imply that the neurons would be unable to respond to the stress caused by intracellular aggregation of Tau and instead the glial cells have elevated levels of stress proteins. In contrast, activation of the innate immune response, as noted by the Unwin group, may reflect activation of microglia and subsequent neuroinflammatory response (Xu et al., 2019). Experiments with cell-type specific read out will ultimately be needed to decide between the three possibilities.

### Stress response

HSP70, BAG3 and the sHSPs have in common that they are all under HSF-1 regulation (Bjork and Sistonen, 2010; Lindquist, 1986; Mathew and Morimoto, 1998). These three components can form a chaperone-complex in the autophagy-pathway, targeting their substrates towards lysosomes for degradation (Stürner and Behl, 2017). A decrease in HSP90 can contribute to the upregulation of HSF-1, as inhibition of HSP90 triggers the heat shock response (Zou et al., 1998). Upon upregulation of HSF-1, there is an elongated occupation of HSF-1 on the HSP70 gene resulting in increased levels of HSP70 (Do et al., 2015; Shapiro et al., 2015). Downregulation of HSP90 and upregulation of sHSP-HSP70-BAG3 may thus be functionally linked in Alzheimer neurons.

### Degradation

Besides activation of HSF-1, a decrease in HSP90 may also have a direct effect on Tau turnover. Inhibition of HSP90 promotes proteasomal degradation of monomeric phosphorylated Tau (Dickey et al., 2007). If a decrease in monomeric Tau degradation is the result of decreased levels of HSP90, this may lead to an accumulation of phosphorylated monomeric Tau, which can give rise to fibril formation. Proteasomal degradation of oligomeric or fibrillar Tau however may not occur, though, as the narrow pore of the proteasome restricts the maximum size of the substrate to be degraded, precluding degradation of protein oligomers or aggregates (Williams et al., 2006).

The other important degradation pathway in AD is the autophagy-lysosome pathway (Wong and Cuervo, 2010). One of the autophagy pathways - macro-autophagy – is upregulated under normal aging conditions (Gamerdinger et al., 2009). The BAG3/BAG1 ratio increases during aging, which is indicative of sequestering towards the autophagy versus proteasomal degradation. BAG1 and BAG3 are both NEFs for HSP70; the HSP70:BAG1 complex targets substrates for proteasomal degradation (Luders et al., 2000) whereas the HSP70:BAG3 complex in cooperation with sHSPs can sequester aggregates for autophagic degradation (Carra et al., 2008). A further increase of the BAG3:BAG1 ratio in AD brain may reflect an even stronger enhanced autophagic activity.

Interestingly, a strong increase of aggrephagy marker SQSTM-1 as observed in AD (Fig. 8B) is indicative of impaired autophagy. Under normal conditions, SQSTM-1 is, together with cargo, cleared by the lysosomes (Bjorkoy et al., 2009). Possibly these observations in AD brain relate to protein aggregates triggering co-operation of sHSPs, HSP70 and BAG3, thereby enhancing the macro-autophagy pathway. However, an immense influx of protein aggregates may overload the autophagic pathway, which in turn may enhance SQSTM-1 levels to prevent clogging of the system.

### Hen and egg question

One of the most stunning questions is why all paralogues of Hsp90 are decreased in AD brains and thus what is upstream of the Hsp90s that may trigger this. In other words, is the decrease in HSP90 cause or consequence. Here we need to consider that the protein levels of the neurons who suffered most from AD cannot be measured, as these neurons already degenerated. The protein levels we study here represent the average in brain regions that contain neurons close to cell death but possibly a majority may not be in their end-stage yet. Such cells may represent the status quo on the road to derailment.

Altered protein levels in AD brain is a good indication of pathway derailment in AD, but there is not necessarily a stringent relation between levels and functionality of a protein. The activity of ATP-dependent chaperones for instance depends on a network of regulatory co-chaperones. Notably, the levels of the co-chaperone-network is largely untouched in Alzheimer, suggesting that alterations in levels of major chaperones such as HSP70 and HSP90 may indeed have functional consequences (Fig. 11B). We note that this is a correlative reasoning, which asks for functional experiments to establish a causal relationship.

In summary, our meta-analysis provides an extensive overview of players in the PQC network in AD, revealing that the majority of factors does not differ between an AD diseased aged brain and a ‘normal’ control brain. It is the specific downregulation of Hsp90 chaperones and upregulation of a pathway centring on the stress inducible HSP70 in AD brains which may result in insufficient capacity for the affected neurons to deal with protein aggregation. We conclude that the PQC including pathways such as the stress response and autophagy play a key role in derailment in AD, which make them interesting targets to interfere in neurodegeneration.

## ACKNOWLEDGEMENTS

We thank Luca Ferrari, Hannah Verwei and Marijke Stokkel for comments on the manuscript, and Simone Lemeer for advice on interpreting data. S.G.D.R. is supported by Campaign Team Huntington, a ZonMW TOP grant (‘Chaperoning axonal transport in neurodegenerative disease’; No. 91215084) and the Top Sector Chemie (CHEMIE PGT. 2019.008).

## AUTHOR CONTRIBUTIONS

M.B.K and S.G.D.R. conceived the study, M.B.K gathered data, M.B.K. and S.G.D.R. analysed the data, M.B.K made figures, M.B.K and S.G.D.R wrote the manuscript

## METHODS

To give a comprehensive overview of protein expression levels in chaperone networks in AD, we performed a meta-analysis of an online database provided by the Unwin group (Xu et al., 2019). The Alzheimer’s Disease Proteome is a freely accessible database reporting proteomics data of more than 5000 distinct proteins, comparing Alzheimer brains versus age-matched control brains. It provides protein levels of nine AD brains and nine healthy control brains, specifying six brain regions, ranging from mostly unaffected to strongly affected in AD. The hippocampus, cingulate gyrus and entorhinal cortex are brain regions strongly affected in AD, motor cortex and sensory cortex are mildly affected, and the cerebellum remains mostly unaffected. For all distinct proteins the data are based on Low-PH LC-mass spectrometry (Xu et al., 2019). In the database, protein levels are calculated based on peptide presence and analysed according to a Bayesian protein-level differential quantification. The database indicates protein levels as logarithmic fold change value of AD brains compared to the age-matched control brain, averaged for each of the populations. The figures in this paper use the non-logarithmic fold change. A quality indicator for data diversity are False Discovery Rates (FDR). It assesses the percentage of significant false positive hits. An FDR value of 1% indicates a likelihood of a significant false positive finding of 1%, in case of 5% this likelihood is 5%.

We visualised all protein levels in a heat-map format, along a double gradient colour-scheme. Baseline for protein levels (1.0) was set to white and can be considered as 0% change of AD compared to control brain. All increases in protein levels are scaled along a blue gradient, whereas decreases in protein levels are scaled along a red gradient. In case some protein level data is unavailable in specific brain regions, boxes are coloured grey.

